# Phase transition of tensin-1 during the focal adhesion disassembly and cell division

**DOI:** 10.1101/2022.11.16.516818

**Authors:** Yuh-Ru Julie Lee, Soichiro Yamada, Su Hao Lo

## Abstract

Biomolecular condensates are non-membranous structures that are mainly formed through liquid-liquid phase separation. Tensins are focal adhesion (FA) proteins linking the actin cytoskeleton to integrin receptors. Here we report that GFP-tagged tensin-1 (TNS1) proteins at physiological levels phase separate to form biomolecular condensates in TNS1 knockout cells. Live cell imaging showed that new TNS1 condensates are budding from the disassembling ends of FAs, and presence of these condensates is cell cycle dependent. TNS1 condensates dissolve immediately prior to mitosis and rapidly reappear while post-mitotic daughter cells establish new FAs. TNS1 condensates contain selected FA proteins and signaling molecules such as pT308Akt but not pS473Akt, suggesting previously unknown roles of TNS1 condensates in disassembling FAs, as the storage of core FA components and the signaling intermediates.

---

Biomolecular condensates are micron-scale membrane-less subcellular organelles that contain biomolecules such as proteins and nucleic acids (1). They form spherical shapes, may merge into one upon contact, and are able to exchange with the surrounding medium within seconds (1–5). They are mainly organized through liquid-liquid phase separation driven by multivalent molecular interactions (1). Although their functions are not fully understood, formation of biomolecular condensates can dramatically enrich relevant molecules, such as enzymes and their substrates, transcription factors and coactivators, or elongation factors, within a very small volume of the compartment and significantly accelerate the reactions involved (4–6).

The involvement of phase separation associated with focal adhesion (FA) proteins was recently explored. The small GTPase regulators GIT/PIX proteins form condensates *in vitro* and in overexpressing cells (5). These condensates serve as modules that can be recruited via specific binding partners to FAs, neuronal synapses, or cell-cell junctions, enabling spatiotemporal regulation of GTPase activities (5). LIMD1 proteins can form opto-genetically induced condensates in cells and be recruited to nascent FAs in a mechanical force dependent manner to facilitate FA maturation (7). Additionally, p130Cas and FAK undergo phase separation and are sufficient to reconstitute kindlin-dependent integrin clustering *in vitro* (8). These findings have specified important roles of phase separation of FA proteins. The genesis and dynamics of these condensates in live cells are largely unknown. Therefore, the complete picture of condensate functions at FAs remains elusive.

## Results and Discussion

Tensin-1 (TNS1) is a FA protein that links the actin cytoskeleton to integrins and mediates outside-in and inside-out signal transductions that are critical for many biological events (9). Analysis of the human TNS1 protein sequence using the IUPred3 revealed a long stretch intrinsically disordered region that is favorable for forming biomolecular condensates (fig 1A). As shown by live cell imaging of TNS1 knockout MDCK cells (10) expressing GFP-tagged TNS1 at a lower than the endogenous TNS1 level (fig 1B), TNS1 formed spherical structures that fused upon contact (fig 1C, Movie S1) and exchanged with the surrounding medium demonstrated by Fluorescence Recovery after Photobleaching (FRAP) assays (fig 1D&E, Movie S2). These findings indicate that TNS1 proteins at the physiological level form membrane-less biomolecular condensates in cells.

**Figure 1.**
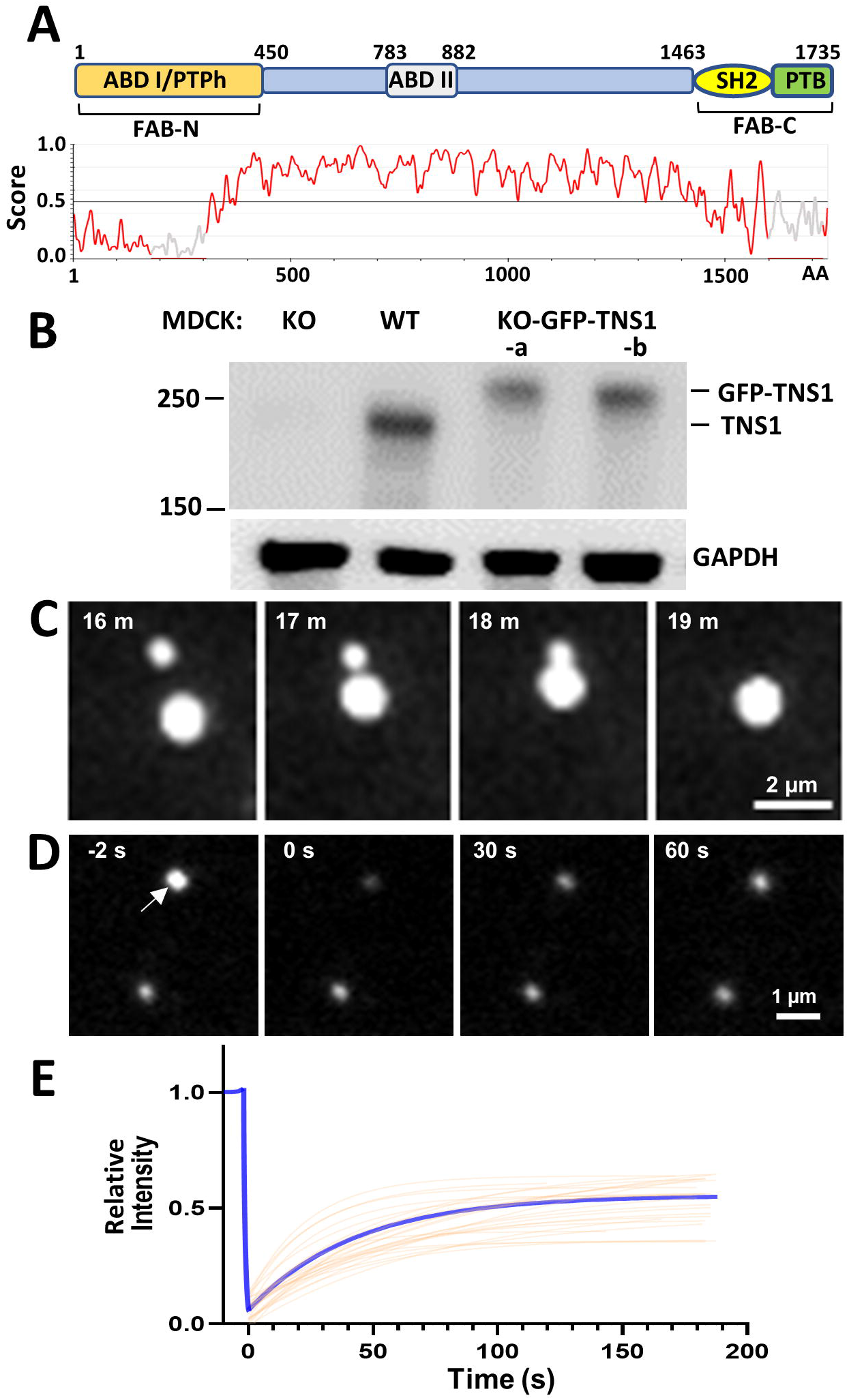
GFP-tagged TNS1 proteins form condensates in cells. (A) Schematic diagram of TNS1 and IUPred3 identification of disordered region of human TNS1 protein. A score > 0.5 indicates disordered. ABD: Actin-Binding Domain, PTPh: Protein Tyrosine Phosphatase homology, SH2: Src Homology 2, PTB; PhosphoTyrosine-Binding, FAB: Focal Adhesion-Binding. (B) Equal amounts of protein lysates from MDCK wildtype, TNS1-knockout, and two TNS1-knockout re-expressing GFP-TNS1 cell lines were immunoblotted against TNS1 and GAPDH antibodies. (C) Live-cell imaging showing two TNS1 condensates were fused into one. (D) Live-cell imaging of FRAP studies. The top TNS1 condensate (arrow) larger than the laser beam was bleached at 0’’ and its fluorescence recovery was recorded in seconds; the bottom punctate was an unbleached control. (E) The average fitting curve (blue) of the FRAP experiments suggested the average half-time of recovery (t1/2) was 30 seconds and the average percentage of recovery was 55 % (n=25).

Interestingly, new TNS1 condensates were budding from rear ends of nascent FA sites. As the nascent FA extended toward leading edge of plasma membrane, the TNS1 signal intensified and assembled into a punctum at the rear end of FAs and segregated away toward the cell center (fig 2A-C, Movie S3). The formation of TNS1 condensates is most obvious in spreading cells without any TNS1 condensates integrating into FAs. These data demonstrate that TNS1 condensates are derived from disassembling FAs. While FA condensates are thought to be precursors to nascent FAs (5, 7, 8), our results support that phase separation of FA proteins may drive FA disassembly.

**Figure 2.**
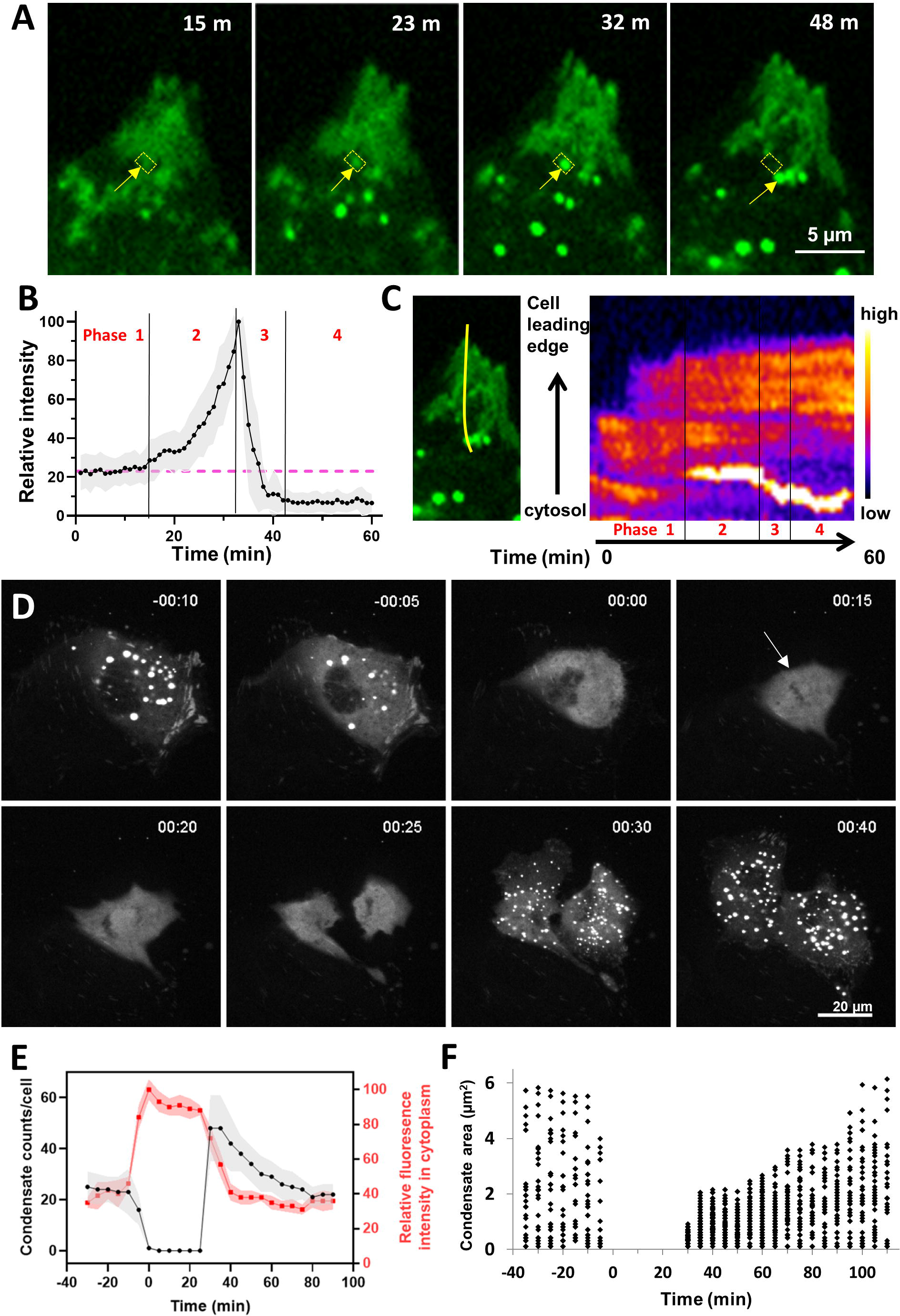
Dynamics of cellular TNS1 condensates. (A) TNS1 condensates (arrows) were derived from and displaced the rear end of GFP-TNS1 positive focal adhesions. (B) Intensity quantification of developing condensates (as framed in yellow square in (A)) revealed four phases: pre-condensate formation (phase 1), condensate developing (phase 2), departing (phase 3) and departed (phase 4). The intensity values were measured and normalized based on the highest intensity during each time course as 100. The averages (connected dots) and standard deviations (shaded area) were plotted (n=20 condensates from 5 cells). The intensities at phase 4 were significantly lower than phase 1 (by a two tailed Student’s t-test, p<0.0001), suggesting that condensate formation/departure has displaced a portion of FA. (C) A kymograph showed the development of condensate in the tracking area (the yellow line). (D) Live cell imaging of GFP-TNS1 condensates during mitosis (hr:min): prophase (00:00, defined by the nuclear envelope breakdown), metaphase (00:15, the GFP excluded area (arrow) indicated chromosome alignment at metaphase plane), cytokinesis (00:25, indicated by the formation of two daughter cells). No condensates were detected from 00:00 to 00:25. (E) The condensate counts in each cell (left vertical axis) and the relative fluorescence intensity in cytoplasm (right vertical axis, the highest intensity was set as 100). Averages were connected by lines and shaded regions denote the standard deviations along the time course (n=6 dividing cells); (F) The area/size of condensates (n=15 to 100 each time point).

Intriguingly, TNS1 condensates dissolved when cells undergo mitosis (fig 2D, Movie S4). Condensates started to disappear minutes prior to prophase, and no condensate was detected from prophase to cytokinesis (fig 2D&E). Parallel to the disappearance, the fluorescence intensities in cytoplasm were markedly increased (fig 2D&E), suggesting that TNS1 condensates were dissolved, instead of degraded, into the cytoplasm. Although trypsin induced cell detachment and spherical cell morphology similar to mitotic cells, the TNS1 condensates remained in the cytoplasm (Movie S5), indicating that the dissolution process is not dependent on attachment status or cell shape, but rather cell cycle regulated. The current literature has focused on the assembly of biomolecular condensates and little is known about the disassembly of biomolecular condensates. Our results indicate that TNS1 condensate disassembly is rapid and tightly regulated by a currently unknown mechanism.

Upon the completion of cytokinesis, many smaller TNS1 condensates rapidly appeared (fig 2D-F). The reappearance of TNS1 condensates corresponded with re-attaching/flattening of daughter cells after cell division, a process requiring rapid formation and turnover of nascent FAs. Since the number of TNS1 condensates increases in post mitotic cells (fig 2F), our observation supports the notion that at least some of TNS1 condensates are spawn from dynamic FAs as a product of FA disassembly rather than consumed as a precursor of FA assembly. After cell division, the average condensate sizes gradually increased, while the condensate numbers reduced, and both values reached pre-mitosis values in 80-90 minutes (fig 2E&F). These findings are consistent with condensates merged by contact-induced fusion (fig 1C).

Since condensates were derived from FAs, we tested and found that FA proteins including ILK, paxillin, ACTN4, DLC1, vinculin, and p130Cas were colocalized to TNS1 condensates. Meanwhile, FAK, zyxin, talin-1, talin-2, ACTN1, VASP, filamin A, Src, and integrin β1 were not (fig 3A). Surprisingly, Akt, a protein ser/thr kinase that mediates many signaling events (11), was detected at TNS1 condensates. T308 and S473 are two major phosphorylation sites on Akt. Phosphorylation of T308 activates AKT kinase activity, and further S473 phosphorylation maximizes Akt’s kinase activity (11, 12). Interestingly, only pT308 of Akt was detected in TNS1 condensates (fig 3B). These findings indicate that TNS1 condensates selectively recruit partners to form biomolecular condensates that are not identical to GIT/PIX, LIMD1, p130Cas, or FAK condensates (5, 7, 8).

**Figure 3.**
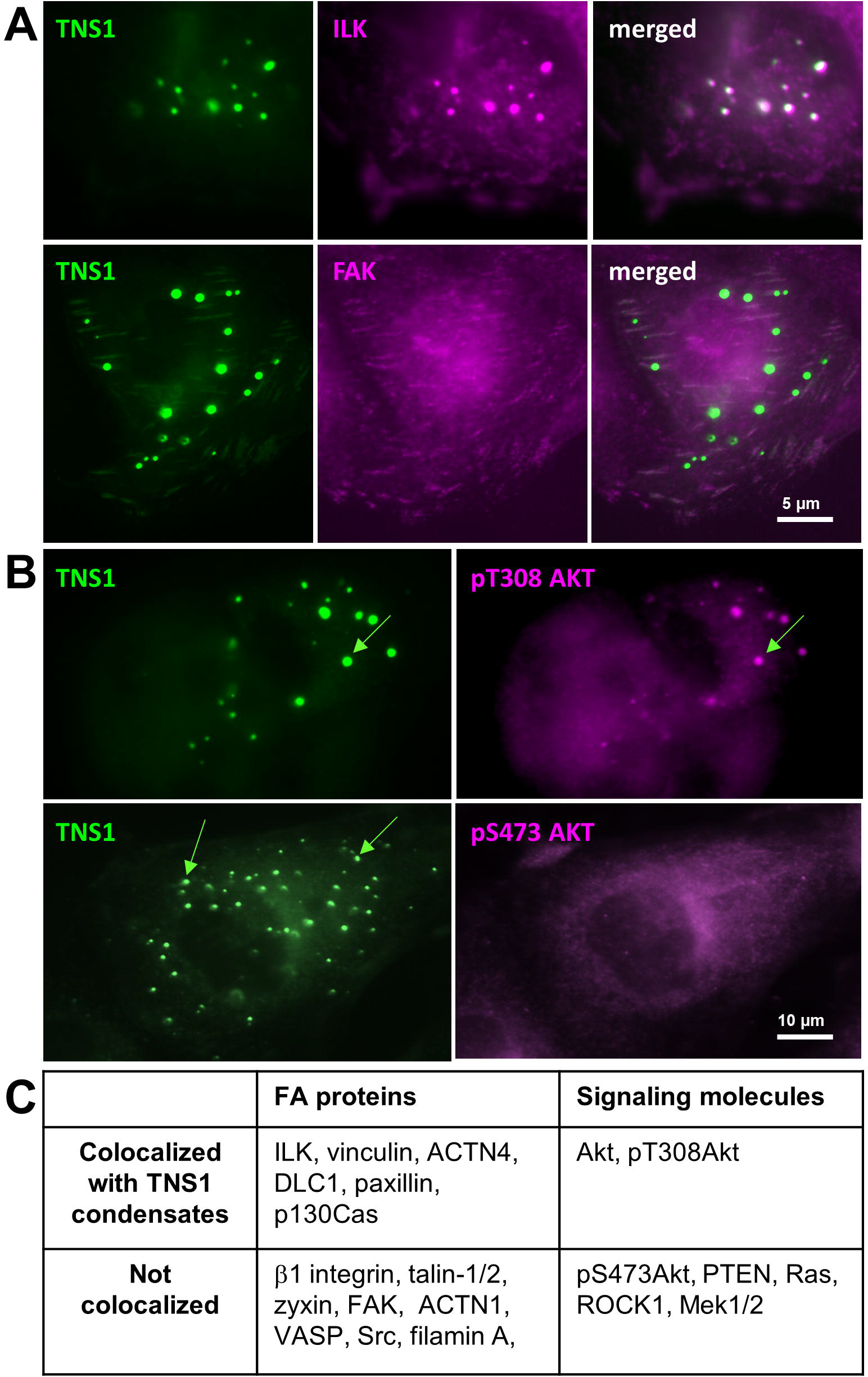
Phase separation of GFP-tagged TNS1 recruits selected proteins to condensates. Cells were fixed and immunofluorescence stained with indicated antibodies. Representative results show that TNS1 condensates colocalized with (A) focal adhesion proteins ILK, but not FAK, or (B) signaling molecules pT308Akt, but not pS473Akt. (C) Summary of colocalization studies.

Based on our findings on TNS1 condensates, we propose that phase separation at FAs promotes FA disassembly and releases biomolecular condensates driven by various FA scaffold proteins, such as TNS1, GIT/PIX, and LIMD1. These condensates store their own sets of FA proteins and/or signaling molecules as future building materials and signaling intermediates in responding to immediate needs of forming new FAs and transmitting signaling cues, respectively. Why TNS1 condensates dissolve prior to cell division is currently unknown, but it is likely that the dissolution allows equal separation of molecules within condensates to daughter cells for re-assembling nascent FAs and/or the dissolution may release key regulators that are critical for processing mitosis. Our current studies open new areas of research on the dynamics, regulations, and function of biomolecular condensates.

## Materials and Methods

Details of cell culture, live cell imaging, FRAP, kymograph, and immunofluorescence staining are provided in *SI*.

## Acknowledgments

We thank Po-Yuan Tung and Jordan Lang for their efforts on the early stage of this work. This work was supported in part by the Team Research Award from the Department of Biochemistry and Molecular Medicine, UC Davis.

