## Supporting information for "Phase transition of tensin-1 during the focal adhesion disassembly and cell division"

**Videos**:

Supplement movie S1 (for Fig 1C). MDCK^Tns1KO^-GFP-TNS1 cells were recorded in time lapse (hr:min). Two condensates (arrows) were fused into one. Scale bar, 10 µm.

Supplement movie 2 (for Fig 1D). FRAP of TNS1 condensates in MDCK^Tns1KO^-GFP-TNS1 cells. The top GFP-TNS1 condensate (yellow box) was bleached at 00:00 (min:sec) and its fluorescence recovery were recorded in a time course; the bottom condensate was an unbleached control. Scale bar, 1 µm.

Supplement movie S3 (for Fig 2A). Development of TNS1 condensates at focal adhesion sites. MDCK^Tns1KO^-GFP-TNS1 cells were recorded in time lapse (hr:min). GFP-TNS1 condensates (arrow) were budding from GFP-TNS1 positive focal adhesion sites. Scale bar, 10 µm.

Supplement movie S4 (for Fig 2D). Presence of TNS1 condensates is cell cycle dependent. MDCK^Tns1KO^-GFP-TNS1 cells were recorded in time lapse (hr:min). Scale bar, 20 µm.

Supplement movie S5. TNS1 condensates were present after the cell was rounded up by trypsin treatment. Time stamp in hr:min; scale bar, 10 µm.

**Extended methods:**

**Cell culture -** MDCK^Tns1KO^-GFP-TNS1 cells were generated by transfection of pEGFP-TNS1 plasmids into MDCK^Tns1KO^ cells (10) and by selection of G418 resistant clones that expressed GFP-TNS1 proteins similar to or lower than endogenous TNS1 protein levels in MDCK cells.

**Live cell imaging** - Cells were grown on glass bottom dishes and live images were acquired as described (13) using a Zeiss AxioObserver equipped with a Yokogawa CSU-10 spinning disk confocal system, a Photometrics CoolSNAPHQ2 camera, and the Slidebook software (Intelligent Imaging Innovations). Images and data were analyzed using Fiji, Microsoft Excel, and the GraphPad Prism Software. The “Multi Kymograph” function in Fiji was used to kymographs. To measure the condensate area and counts in each cell, images were first converted to binary in the threshold range above intensity of FAs and the highest intensity within images, and then were analyzed with the particle analysis function in ImageJ.

**Fluorescence recovery after photobleaching** - The fluorescence recovery was recorded every 2 seconds for 3 minutes as described (13). An ImageJ macro was used to track condensates and measure their corresponding fluorescence intensities. The half-time (t1/2) and the recovery percentages were calculated by fitting the data with non-linear regression (GraphPad Prism Software).

**Immunofluorescence staining**- Cells grown on collagen-coated coverslips were fixed. After incubated with indicated first antibody, coverslips were washed with PBS and incubated with rhodamine-conjugated 2^nd^ antibody. Images were taken using the AxioObserver confocal microscope.

**Antibodies-** Antibodies against ILK (#3862), ACTN1 (#6487), FAK (#13009), Src (#2123), Akt (#4691), pT308Akt (#13842), pS473Akt (#4060), PTEN (#9188), Ras (#8955), Mek1/2 (#8727), ROCK1 (#4035), β-catenin (#8480) were from Cell Signaling Technology. Antibodies against DLC1(#612021), p130Cas (#610271), zyxin (#3610521), VASP (#610448), β1 integrin (#553715) were from BD Biosciences. Talin-1 (MCA4770) and Talin-2 (MCA4771) antibodies were from Bio-Rad. Antibodies against Vinculin (V9131, Sigma), Filamen A (MAB1680) antibodies were from Millipore Sigma. ACTN4 (42-1400) antibody was from Invitrogen. Paxillin (B7192) antibody was from Assay Biotech. TNS1 antibody was made in Lo’s lab.
